# Effects of the exercise-inducible myokine irisin on proliferation and malignant properties of ovarian cancer cells through the HIF-1 α signaling pathway

**DOI:** 10.1101/2021.08.18.456863

**Authors:** Marziyeh Alizadeh Zarei, Elahe Seyed Hosseini, Hamed Haddad Kashani, Ejaz Ahmad, Hossein Nikzad

## Abstract

**Background:** Exercise has been shown to be associated with reduced risk and improving outcomes of several types of cancers. Irisin −a novel exercise-related myokine- has been proposed to exert beneficial effects in metabolic disorders including cancer. No previous studies have investigated whether irisin may regulate malignant characteristics of ovarian cell lines.

**Methods:** In the present study, we aimed to explore the effect of irisin on viability and proliferation of ovarian cancer cells which was examined by MTT assay. Then, we evaluated migratory and invasive ability of the cells via transwell assays. Moreover, the percentage of apoptosis induction was determined by flowcytometery. Furthermore, the mRNA expression level of genes related to the aerobic respiration (HIF-1α, c-Myc, LDHA, PDK1 and VEGF) were detected by real-time PCR.

**Results:** Our data revealed that irisin treatment significantly attenuated the proliferation, migration and invasion of ovarian cancer cells. Besides, irisin induced apoptosis in ovarian cancer cells. We also observed that irisin regulated the expression of genes involved in aerobic respiration of ovarian cancer cells.

**Conclusion:** Our results indicate that irisin may play a crucial role in inhibition of cell growth and malignant characteristics of ovarian cancer. This findings may open up avenues for future studies to identify the further therapeutic use of irisin in ovarian cancer management.

## 1. Introduction

Ovarian cancer (OC), as the most common and lethal gynecologic malignancies, is the fifth cause of cancer-related death in women globally [1, 2]. Due to the fact that OC is usually asymptomatic at early stages, most of the patients will ultimately progress to advance stages, resulting in a high rate of mortality [3, 4]. Despite recent advances, the lack of effective screening methods and therapeutic modalities are the major hurdles to improve prognoses for management of the disease and the main cause of 5-year survival rate of less than 40% [5]. Thus, exploring novel avenues to improve the status quo is urgently needed.

Obesity has been widely reported to be associated with the metabolic diseases including cancer [6–8]. Notably, obese people are usually at increased risk of developing cancers due to the higher levels of insulin and insulin-like growth factor-1, which may be responsible for certain tumors promotion [9]. Exercise is one of the several modifiable factors known to reduce the risk of developing human malignancies [10–13]. It has been also well documented that exercise has many benefits after diagnosis. There is increasingly mounting evidence that exercise relieve many of the common side effects contributed to the conventional cancer therapy among patients, resulting in a better overall quality of life [14–17]. Besides, a number of observational cohort studies support the view that cancer survivors who had a regular exercise suffered from a lower risk of relapse or cancer-related death after starting treatment regimens [18–20].

Recently, skeletal muscle is gaining special interest in terms of releasing various myokines which exert beneficial effects in metabolic disorders. Of these, irisin, a novel promising identified myokine releasing from skeletal muscle following exercise, is believed as a potential therapeutic in a variety of diseases [21, 22]. Previous studies have documented the positive association between irisin and body energy expenditure as well as insulin sensitivity [23]. Additionally, irisin has been verified to have a pivotal role in smooth muscle cell phenotype modulation by regulating endothelial cell proliferation, apoptosis and migration [24]. Due to the strong metabolic effects of irisin on several tissue types, it is questionable whether irisin has the ability to modulate malignant features of cells and tissues similar to other myokines [25]. In the past decade, the relationship between irisin and different cancers have been the focus of many studies. Represented data from a recent study found a lower level of serum irisin in patients with breast cancer, while other studies indicated that irisin was significantly increased in gastrointestinal cancer tissues [26–28]. Moreover, a number of studies have detected increased irisin immunoreactivity in ovarian, cervix and breast cancer tissues as well as endometrial hyperplasias [29].

In the context of cancer metabolism, reprogramming of glucose metabolism is a pervasive microenvironmental event in tumorigenesis. To this end, cancer cells preferentially switch from oxidative phosphorylation (OXPHOS) to glycolysis, even under normal oxygen concentration. This particular metabolic profile – defined as Warburg effect- is a key hallmark of many solid tumors including OC [30].

Growing evidence demonstrated that many signal molecules including tumor suppressor genes and oncogenes play substantial roles in conferring metabolic advantages and adaptation to the tumorigenic microenvironment [30]. Of these documented genes, the hypoxia-inducible factor-1α (HIF-1α) is a transcription factor regulating many pivotal pathways in cancerous cells [31].

Although the importance of HIF-1α in the induction of the Warburg effect is largely highlighted, the involvement and influences of other factors should not be underestimated. A number of studies notably suggested that crosstalk between HIF-1α and oncogenic c-Myc lead to increased uptake of glucose and its conversion to lactate which subsequently modulate the cancer cell’s microenvironment through regulation of common downstream target genes, such as pyruvate dehydrogenase kinase 1(PDK1) and lactate dehydrogenase A (LDHA) [32]. It is also noticeable that angiogenesis is necessary for tumor growth and metastasis .HIF-1α also upregulates the expression of the angiogenic growth factors such as vascular endothelial growth factor (VEGF), which in turn stimulates neovascularization a fundamental process for tumor progression [33]. Considering the above mentioned points, we intended to evaluate the effect of irisin as a promising myokine on tumorigenic features of ovarian cancer cells and to regulate genes correlated with aerobic metabolism.

## 2. Materials and Methods

### 2.1. Cell culture and reagents

The ovarian cancer cell lines OVCAR3, SKOV3 and Caov4 were purchased from the national Cell bank of the Pasteur Institute, Tehran, Iran, were cultured in RPMI1640 ( Gibco, Invitrogen corporation) supplemented with10% fetal bovine serum (FBS) and 1% penicillin/streptomycin (100 units/ mL) and maintained under standard conditions (37º C and 5% CO2). Cells were treated with human recombinant irisin (Cayman Chemical CAS NO: 9037-90-5) at different concentrations ranging from 5 to 70 nM dissolved in phosphate buffer saline (PBS).

### 2.2. Proliferation assay

3-(4,5-Dimethylthiazol-2-yl)-2,5-diphenyltetrazolium bromide (MTT) assay was carried out to determine optimum concentration and time course of action of irisin in OC cell lines (OVCAR3, SKOV3 and Caov4). The mentioned cell lines (10,000 cells/well) in 150 μl medium were seeded overnight into 96-well plates and treated with different doses of irisin (5-70 nM) for different time periods (24,48,72, 96 and 120h). Subsequently, 20 μl of MTT solution (5 mg/mL) was added to each well and incubated for 4 h. Then, 150 μl of dimethylsulfoxide (DMSO) was added to each and incubated for 15 minutes at dark. The absorbance at 570 nm was measured using a microplate reader.

### 2.3. Colony formation assay

OVCAR3, SKOV3 and Caov4 were seeded into a 96-plate at a density of 500 cells/well and treated with 5 and 10 nM of irisin. Untreated cells were considered as controls. All three cell lines were grown for 14 days at standard incubation condition (37º C and 5% CO2). After 14 days of incubation, cells were fixed and stained with 0.5% crystal violet. Colonies including more than 50 cells were attended as surviving cells and colony formation efficiency was calculated as opposed to seeded cells.

### 2.4. Migration and Invasion assays

The in vitro migration and invasion experiments were carried out using a 24-well transwell (BD Bioscience, Bedford,MA). For the invasion test, all cells were serum starved the day before the assay. The lower side of the transwell was filled with RPMI 1640 with 10% FBS, and the transwell pre-coated with 50μl of 1:2 Matrigel/RPMI 1640 (Matrigel, Corning incorporated,USA). All cells (50,000 cells) stimulated with irisin (5 and 10 nM) as well as untreated control cells suspended in serum-free medium and were added to the upper surface of the transwell. After incubation for 48h in 37°C, the medium was completely removed from the top part and non-invaded cells were wiped using cotton swabs. The cell invading to the underside of the transwell were fixed and stained in 0.1% crystal violet. The number of invaded cells through the matrigel was enumerated in 5 randomly fields using an inverted microscope (200×). The migration assay was conducted with a similar method, except that cells were plated into uncoated-matrigel transwells and incubated for 24 h period.

### 2.5. Apoptosis detection

Annexin-V/ PI staining kit (Roche Applied Science, Penzberg, Ger many) was used to analyze apoptosis percentage in OC cells according to the manufacturer’s instruction. Briefly, OVCAR3, SKOV3 and Caov4cells were seeded (200,000 cells/well) into six-well plates and incubated overnight. At 70-80% confluency, cells treated with desired concentration of irisin

(10 nM) and incubated for additional 48h. After 48h, cells were trypsinized and washed with PBS twice. Trypsin-digested cells were centrifuged and cell pellet was resuspended in 100 μL of binding buffer and incubated for 15 minutes at 15°C–25°C. Finally, apoptotic cells were analyzed using FACSCalibur flow cytometer within 1h.

### 2.6. Real-time quantitative polymerase chain reaction (qPCR)

Total RNA was extracted from treated cells using Trizol (Invitrogen, California, USA). Synthesis of Complementary DNA was performed by PrimeScript™ RT reagent kit (Takara Bio Inc., Shiga, Japan), and flowingly used as template for Real-time PCR reaction which was conducted by IQ5 (Biorad, Germany). The changes in mRNA expression of genes of interest were normalized to the levels of the glyceraldehyde-3-phosphate dehydrogenase (GAPDH) as internal reference. Real-time data analysis performed using 2^−ΔΔct^ formula. The sequences of specific primers are shown in Table 1.

**Table 1.**
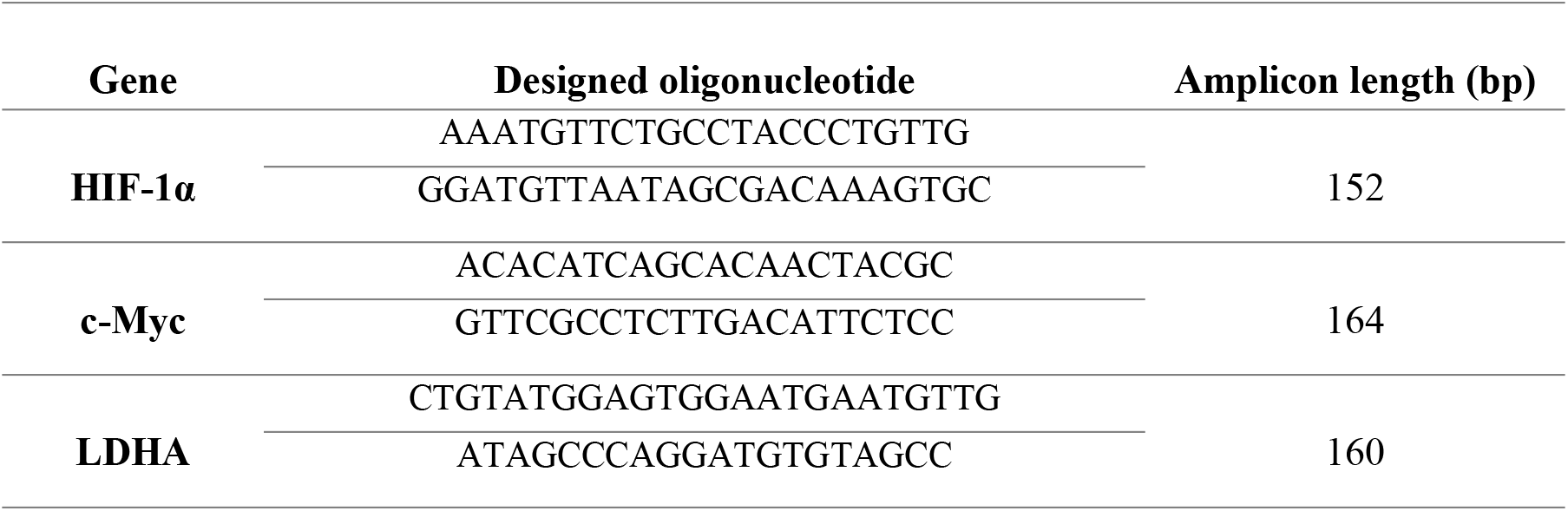
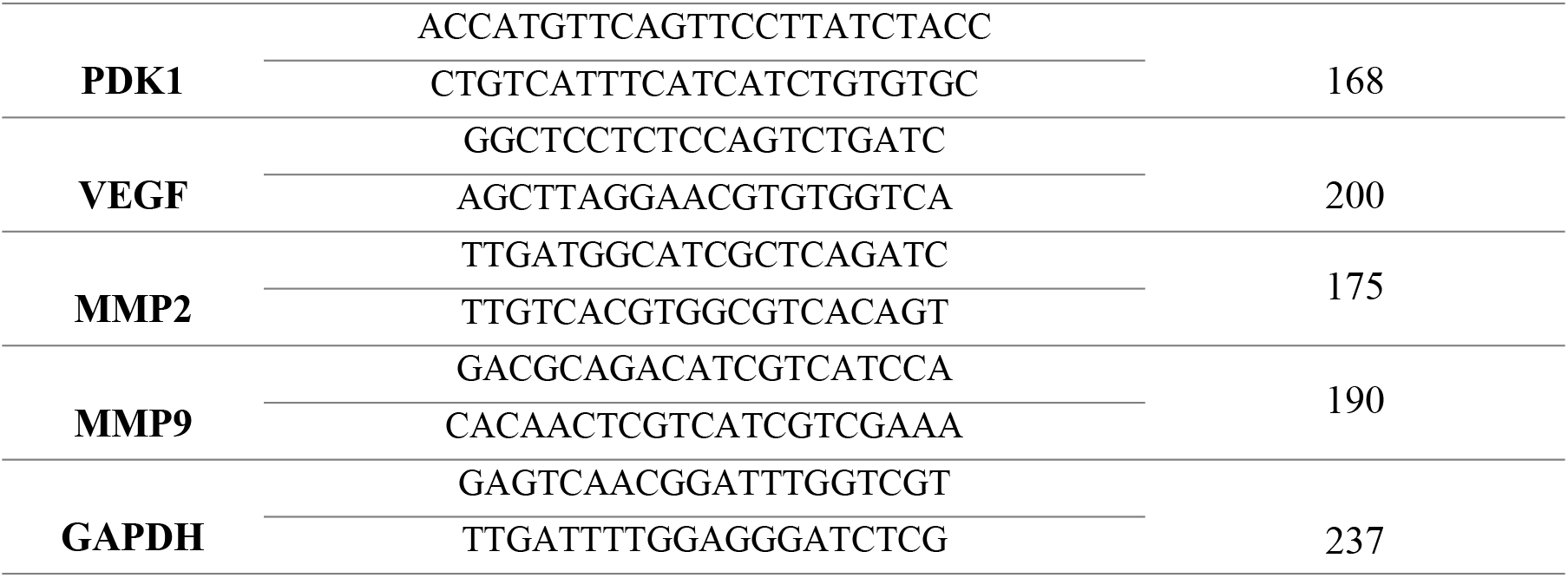
List of specific primers used in real-time polymerase chain reaction assay

### 2.7. Statistical analysis

Statistics were presented in Prism® 5 software (GraphPadSoftware, Inc., La Jolla, CA, USA). The data are expressed as the mean ± standard deviation (SD). All of the experiments repeated at least three times. Comparisons between the groups were conducted using one-way ANOVA analysis of variance followed by Tukey’s HSD multiple comparison test. Statistical significance was considered significant at P < 0.05.

## 3. Results

### 3.1. Regulation of cell proliferation by irisin in ovarian cancer cell lines

MTT assay was used to investigate the effect of various concentrations of irisin within different time periods on OC cells viability. We found no significant changes in the viability of OC cells 24h after treatment. However, our data indicated that irisin significantly suppressed the proliferation of all three cell lines in a time- and dose-dependent manner (Fig. 1).

**Figure1.**
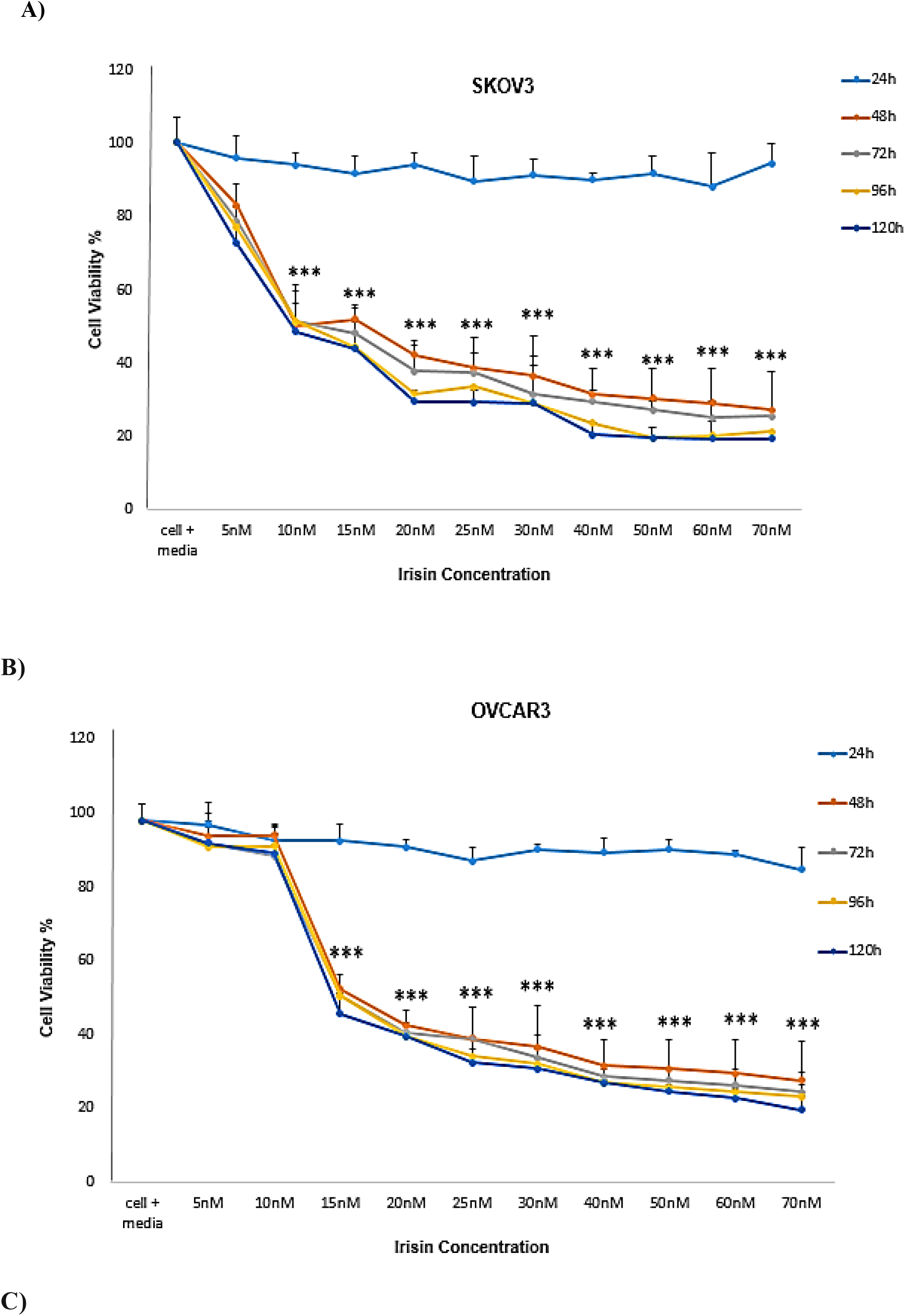
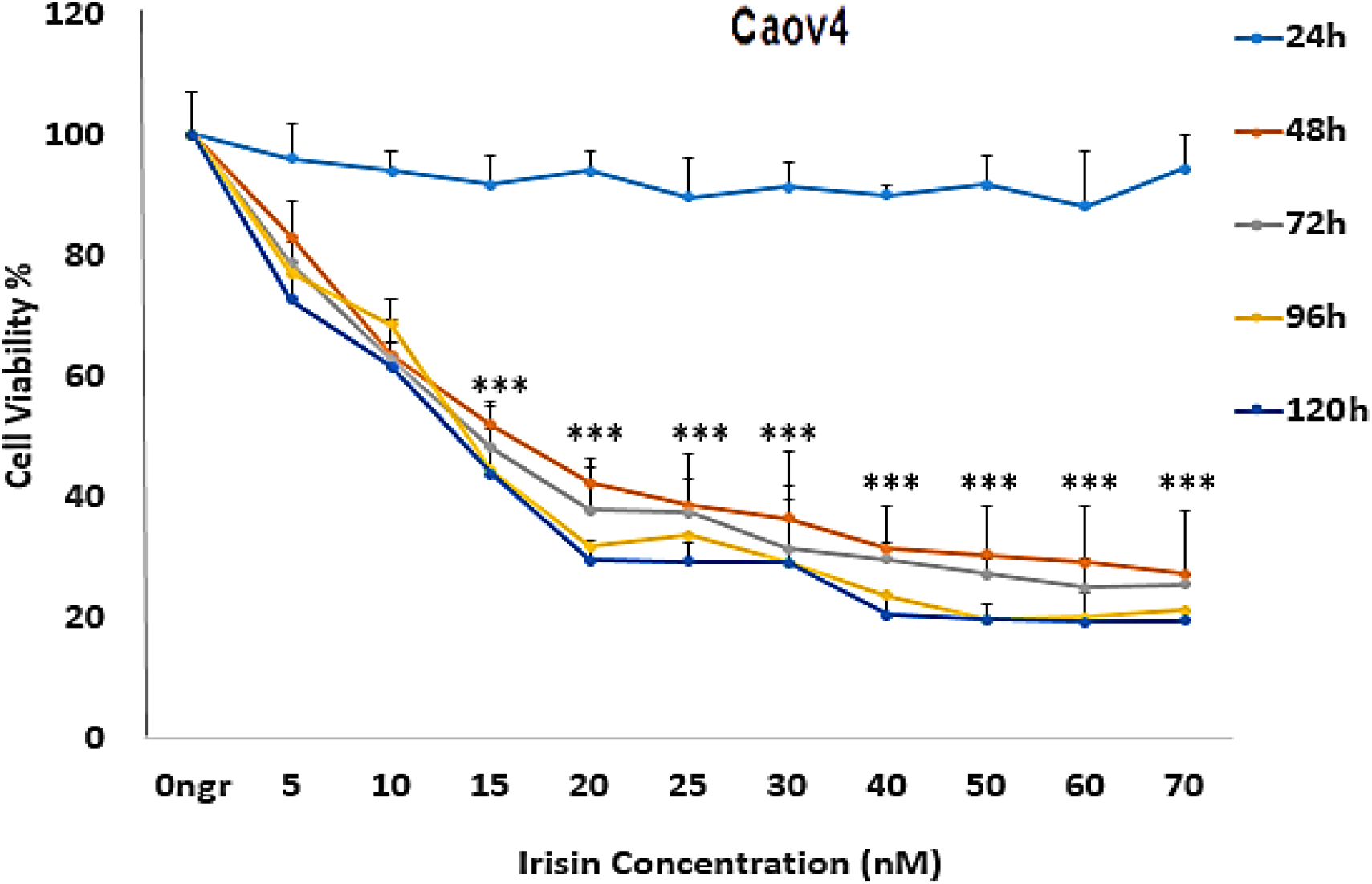
**The** effect of irisin on the proliferation of A) SKOV3 B) OVCAR3 C) Cov4 cells in 24, 48 and 72,96 and 120 hours. **Notes:** Data expressed as mean ± standard deviation; \**P*< 0.05; \*\**P*< 0.01; \*\*\**P*< 0.001 compared to no treated cells

### 3.2. The effects of irisin on colony formation ability of ovarian cancer cell lines

We next evaluated the ability of three cell lines examined to form colonies on 6-well plate plates in the presence or absence of irisin (5 and 10 nm) within 2 weeks. Our findings represented that the colonogenic potential of OC cells was significantly suppressed after treatment with irisin. Although, the greatest inhibitory effect of irisin on cologenic ability was observed in Caov4 cells, the number of colonies formed attenuated in a dose dependent manner in all three cell lines tested, as shown in (Fig. 2).

**Figure2.**
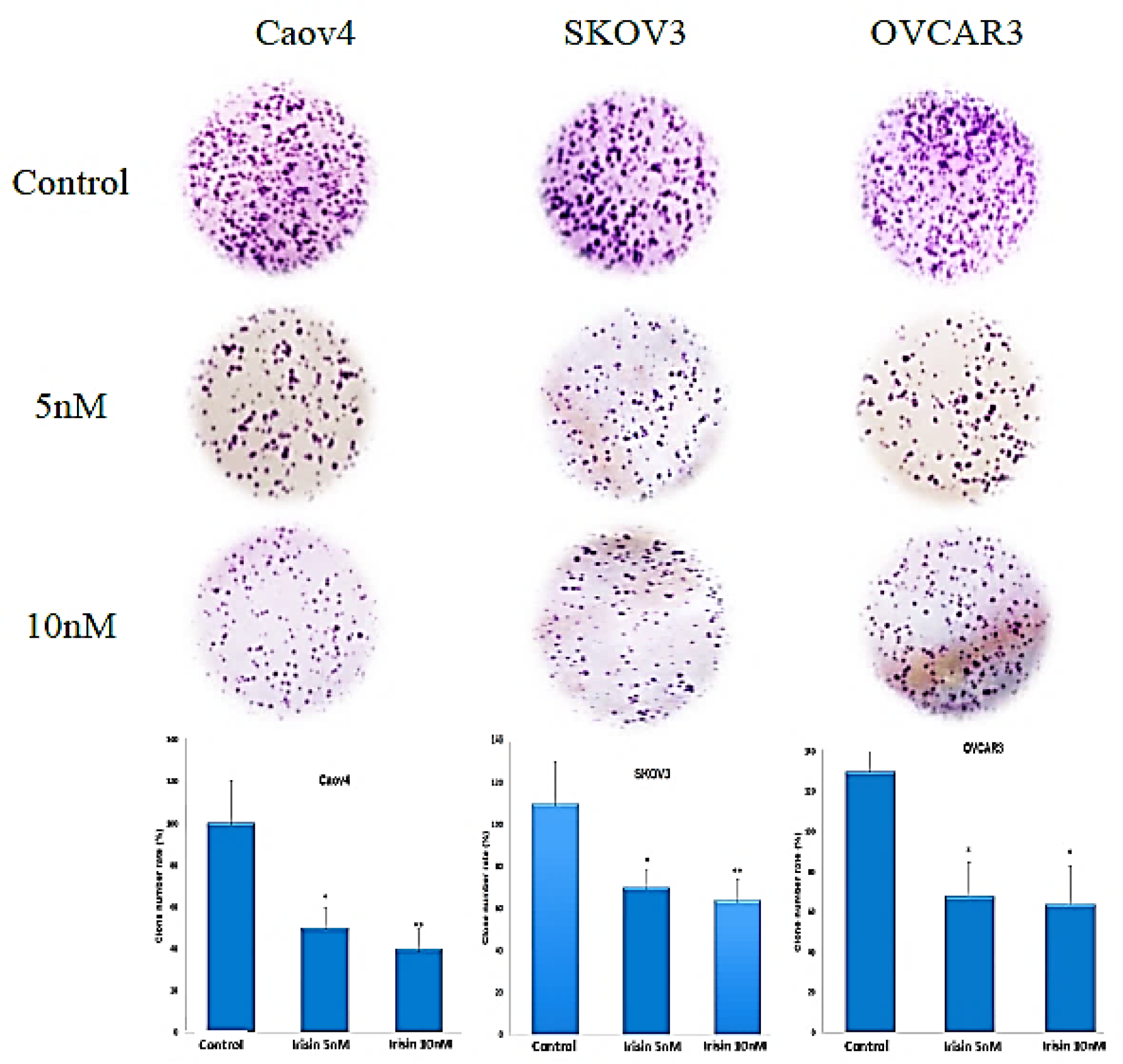
Effect of irisin to inhibit colony formation of Caov4, SKOV3 and OVCAR3 ovarian cancer cell lines. Data shows the representative of three independent experiments. **Notes:** Data expressed as mean ± standard deviation; \**P*< 0.05; \*\**P*< 0.01; \*\*\**P*< 0.001 compared to no treated cells.

### 3.3. The effects of irisin on invasiveness properties of ovarian cancer cell lines

We applied the transwell system to explore the effect of irisin on the migration and invasion of OC cell lines. As our results revealed, irisin effectively reduced the migratory behavior of all three cell lines as compared to control groups. Furthermore, represented data suggested the inhibitory effect of irisin on invasiveness of OC cell lines. The migration and invasion potential of all cell lines subjected to irisin were decreased in a dose-dependent manner (Fig. 3A and 3B).

**Figure3.A.**
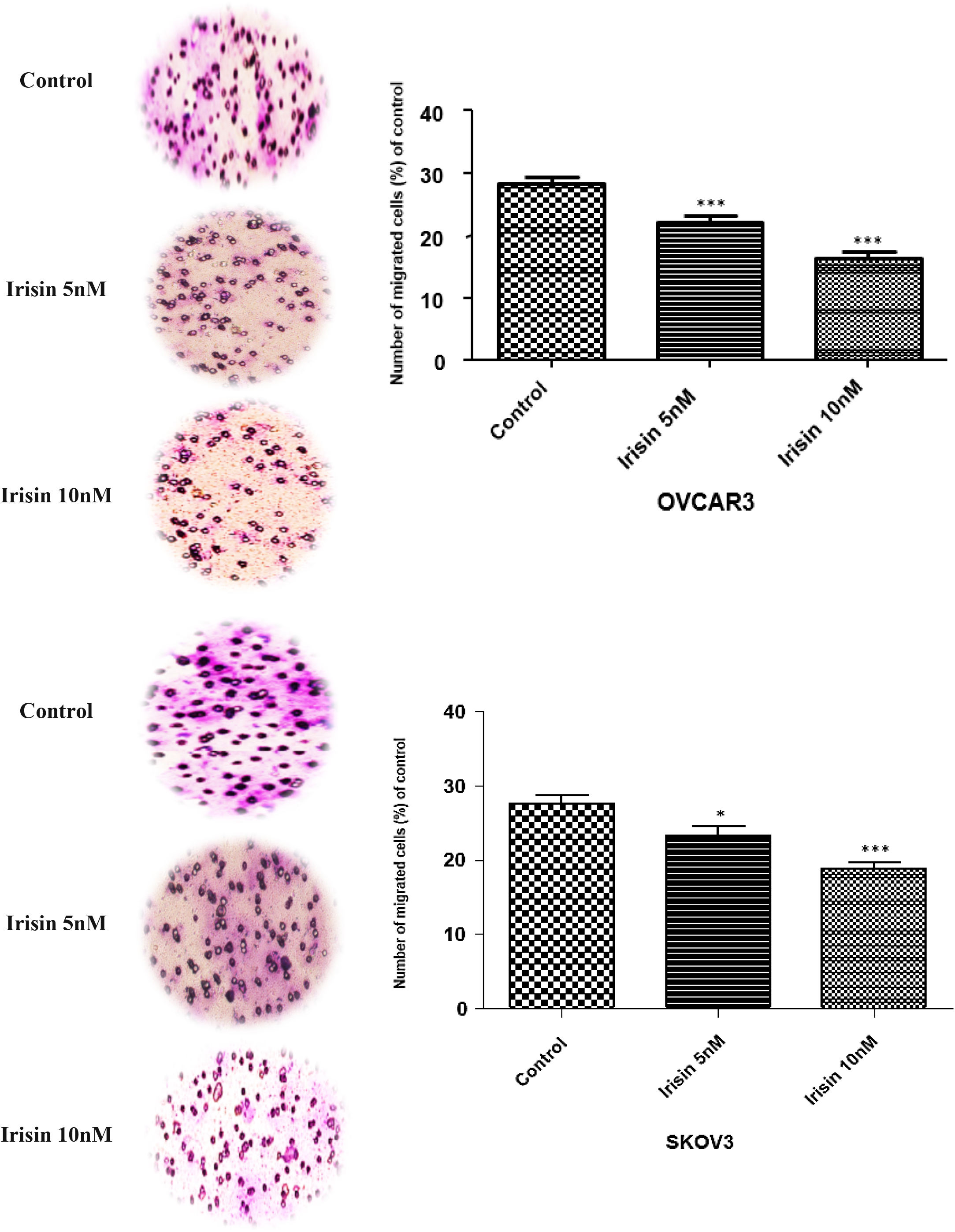
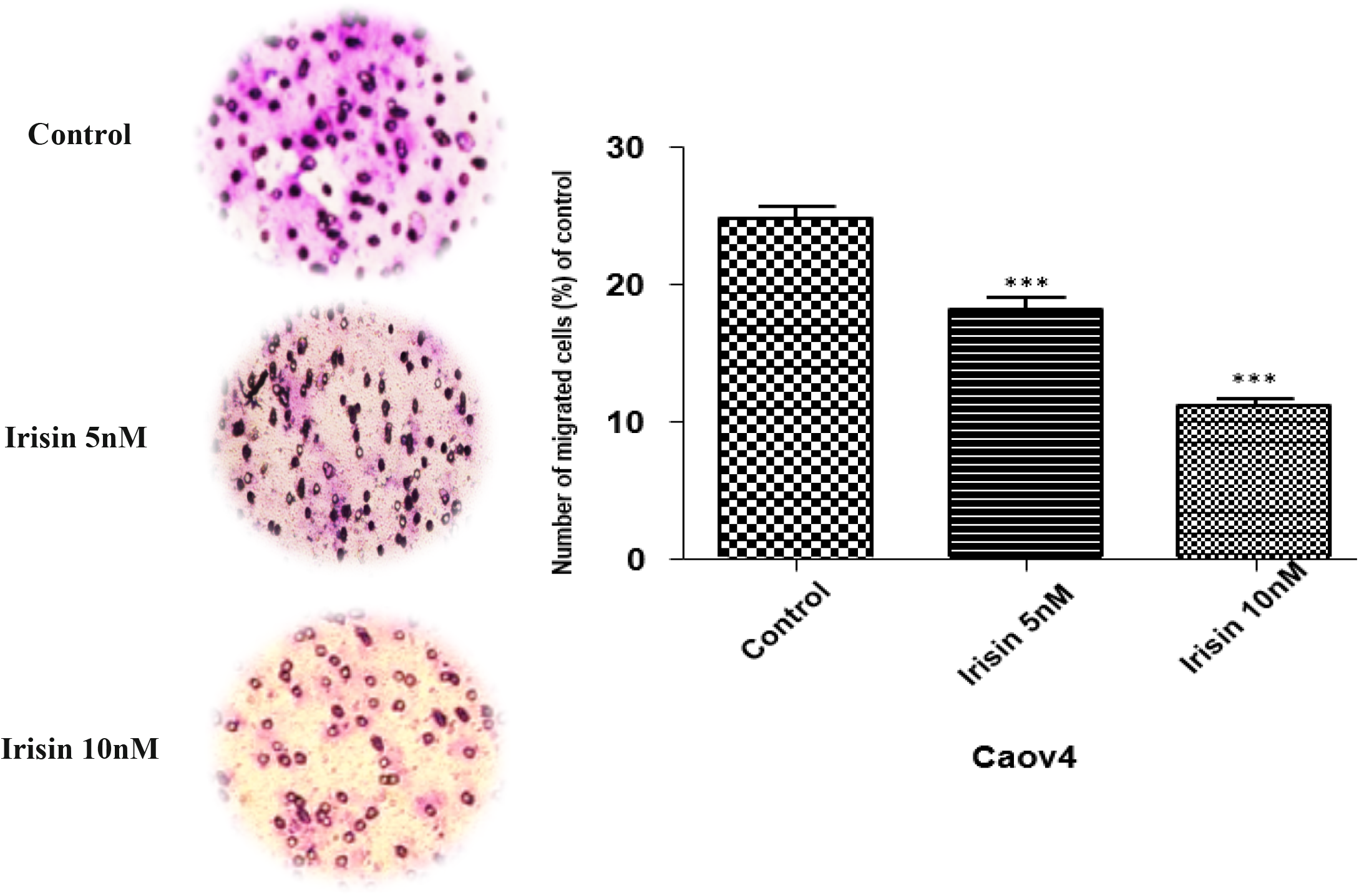
**The effect of** Irisin on the migratory behavior of OVCAR3, SKOV3 and Caov4 cells. **Notes:** Data expressed as mean ± standard deviation; \*\**P*, 0.01; \*\*\**P*, 0.001 compared to control.

**Figure3. B.**
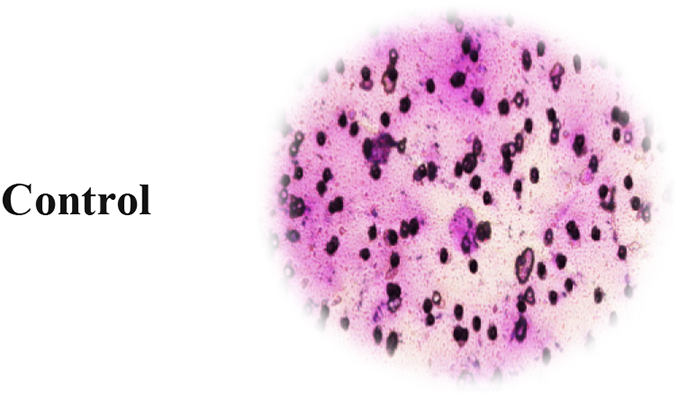
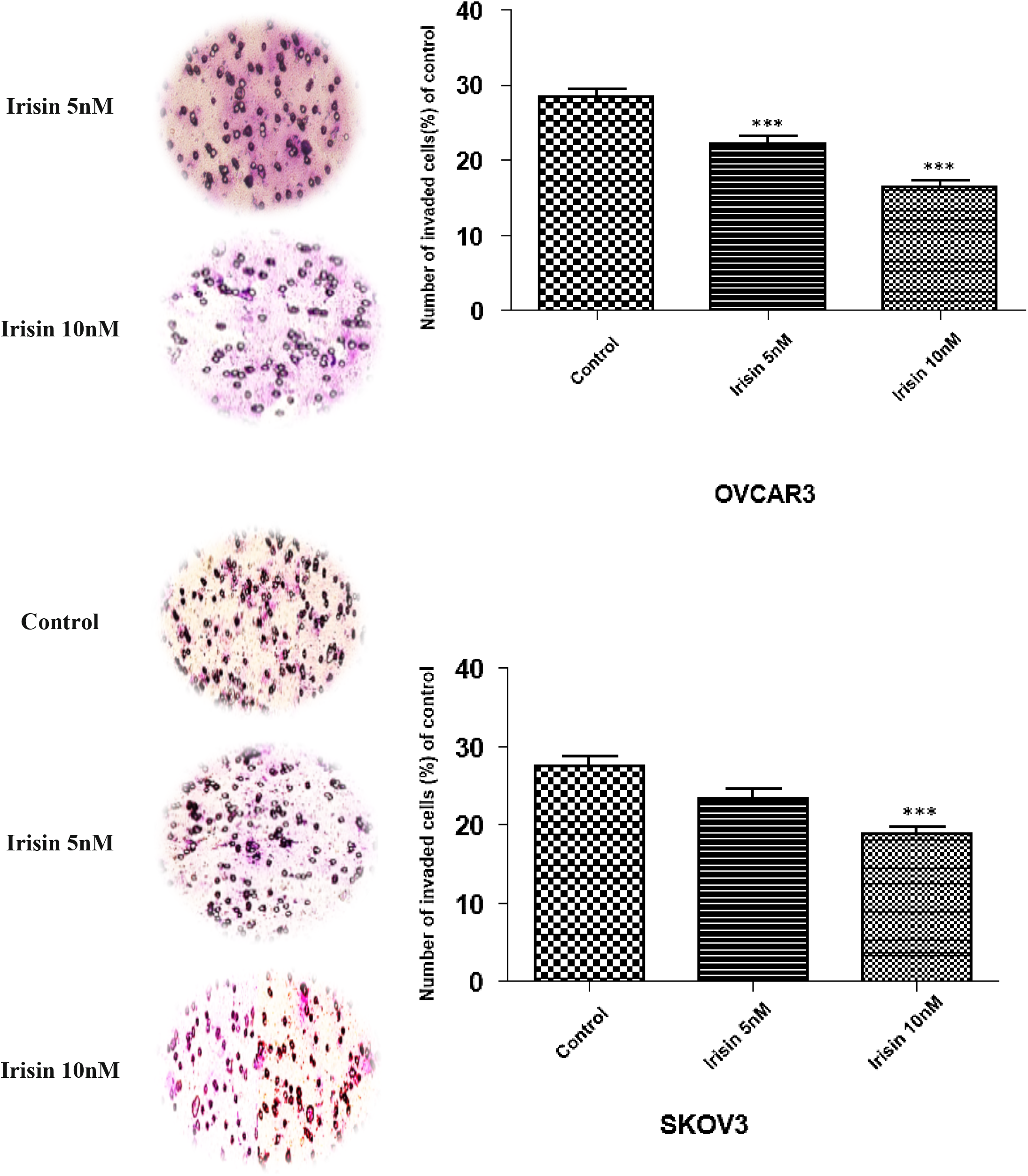
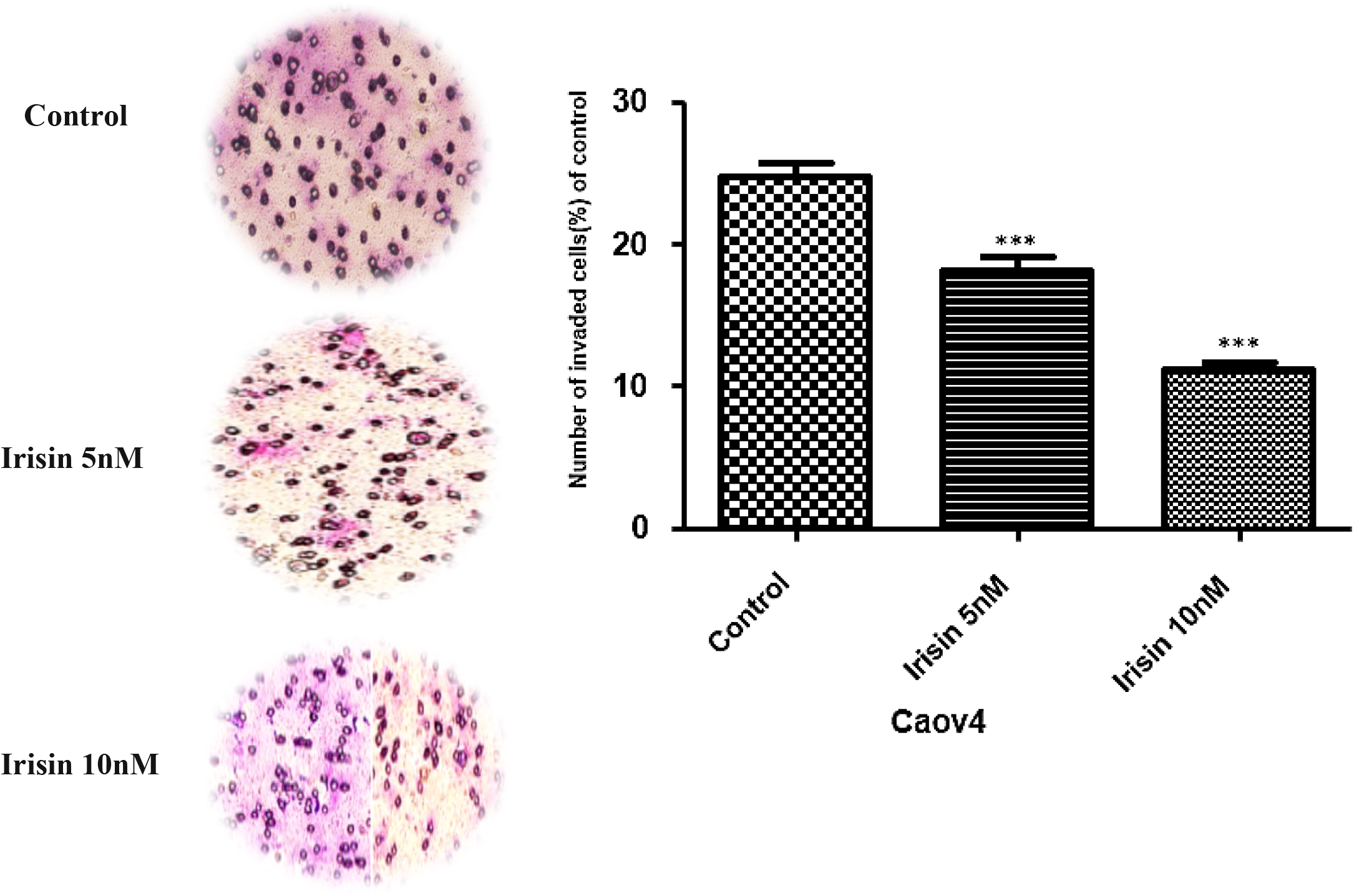
The effect of Irisin on invasiveness of OVCAR3, SKOV3 and Caov4 cells. **Notes:** Data expressed as mean ± standard deviation; \*\**P*, 0.01; \*\*\**P*, 0.001 compared to control.

### 3.4. The effects of irisin on apoptosis induction in ovarian cancer cell lines

Given the inhibitory effects of irisin on OC cell proliferation, we next examined the susceptibility of OC cells to apoptosis in the presence of irisin. Flowcytometry analysis illustrated that irisin caused a significant rise in the percentage of apoptotic Caov4 cells as compared to control cells (P<0.008), however, no significant effect was detected in other two cell lines (Fig.4).

**Figure 4.**
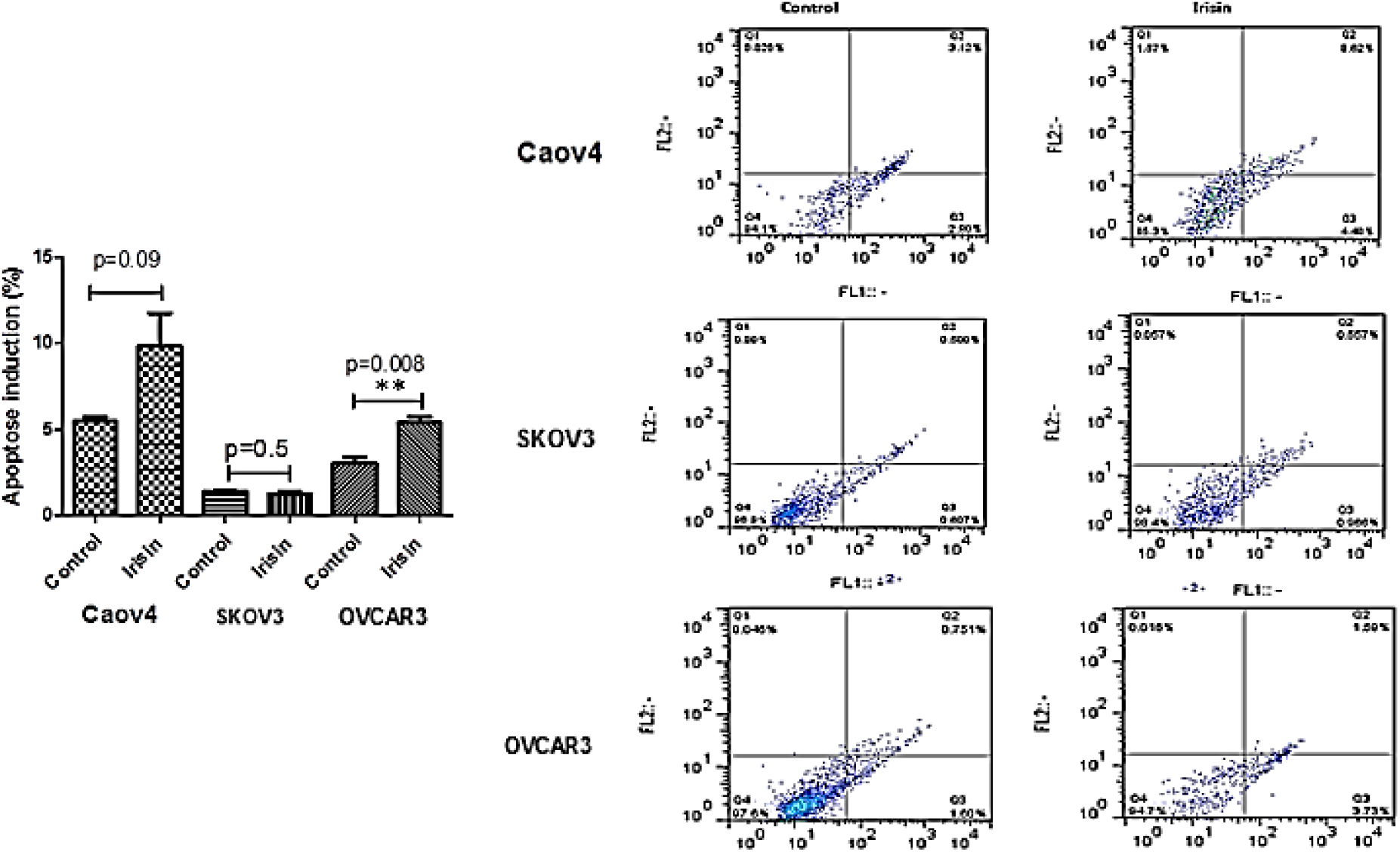
The effect of irisin on apoptosis induction of Caov4, SKOV3 and OVCAR3 in ovarian cancer cells. **Notes:** Data expressed as mean ± standard deviation; \*\**P*, 0.01; \*\*\**P*, 0.001 compared to control.

### 3.5. Irisin affects the expression of aerobic metabolism markers through the HIF1-α signaling pathway in ovarian cancer cells

The expression of aerobic metabolism genes in OVCAR3 and SKOV3 cells in the absence or presence of irisin (5 and 10 nM) was measured using real-time PCR. As depicted in figure 7 A irisin decreased the expression of HIF-1α, C-myc, LDHA and PDK1 genes in SKOV3 cells as compared to control cells, however no significant reduction in the level of mentioned genes was found in OVCAR3 cells. In the case of VEGF expression, irisin attenuated the expression level of this gene in both SKOV3 and OVCAR3 cells. Surprisingly, the greater inhibitory effect of irisin on VEGF expression after treatment with 5 nM of irisin.

### 3.6. The effect of irisin on the expression of MMP2 and MMP9 in ovarian cancer cells

The expression of MMP2 in OC cells treated with irisin was measured using real-time PCR. As depicted in figure 6, irisin remarkably decreased the expression of MMP2 in SKOV3 and OVCAR3 cells as compared to control in a dose-dependent manner. However, irisin had no significant effect on MMP9 reduction in both cell lines.

**Figure 5.**
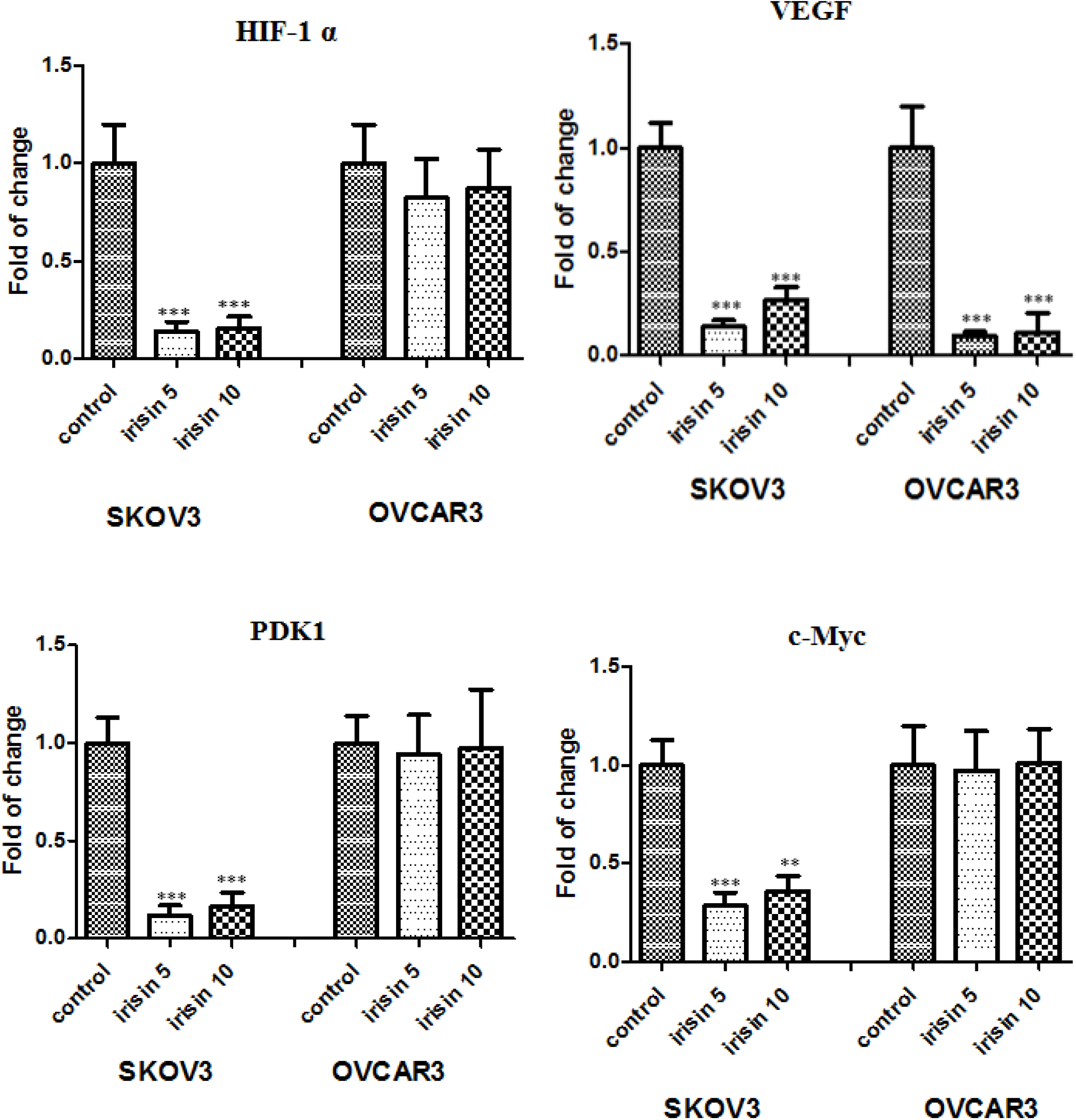
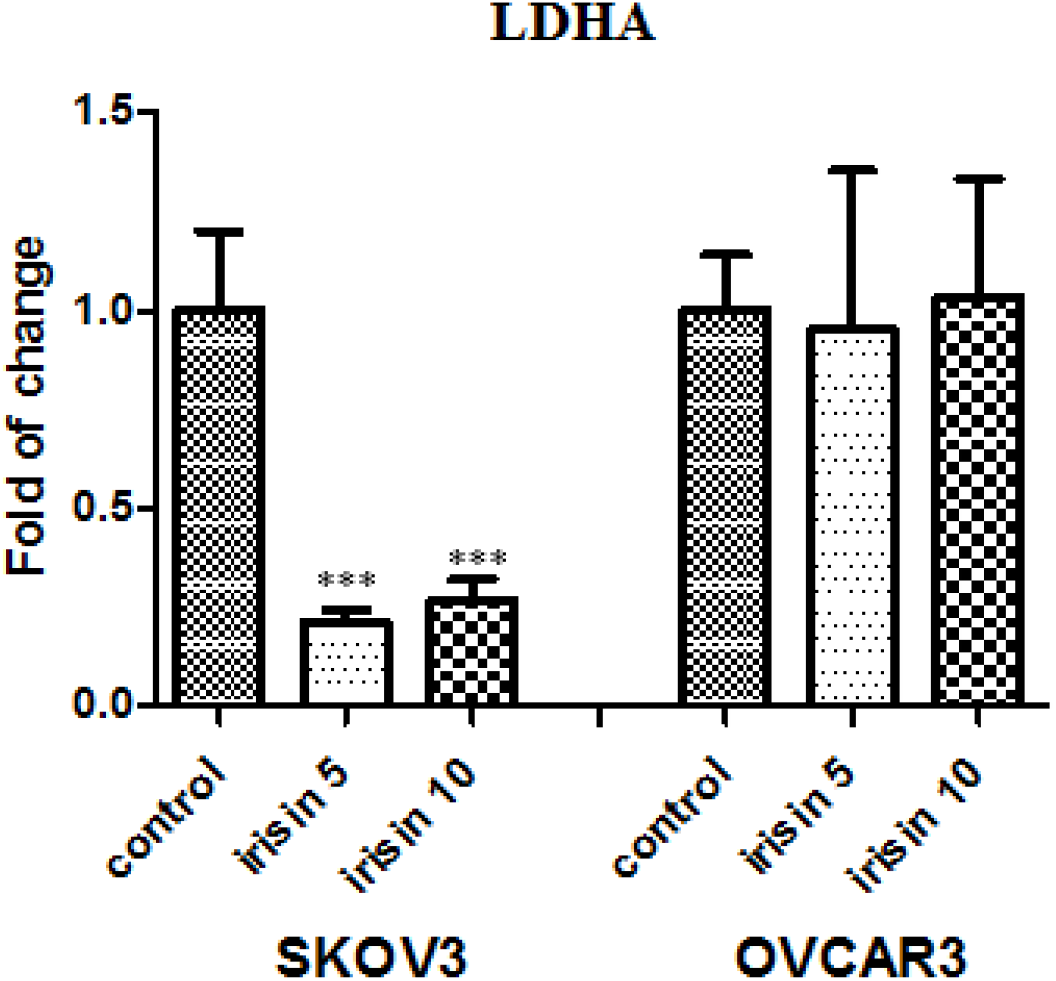
Effect of Irisin on HIF-1 α signaling pathway genes 48h after treatment with 5nM and 10nM of irisin. **Notes:** Data expressed as mean ± standard deviation; \*\**P*, 0.01; \*\*\**P*, 0.001 compared to control.

**Figure 6.**
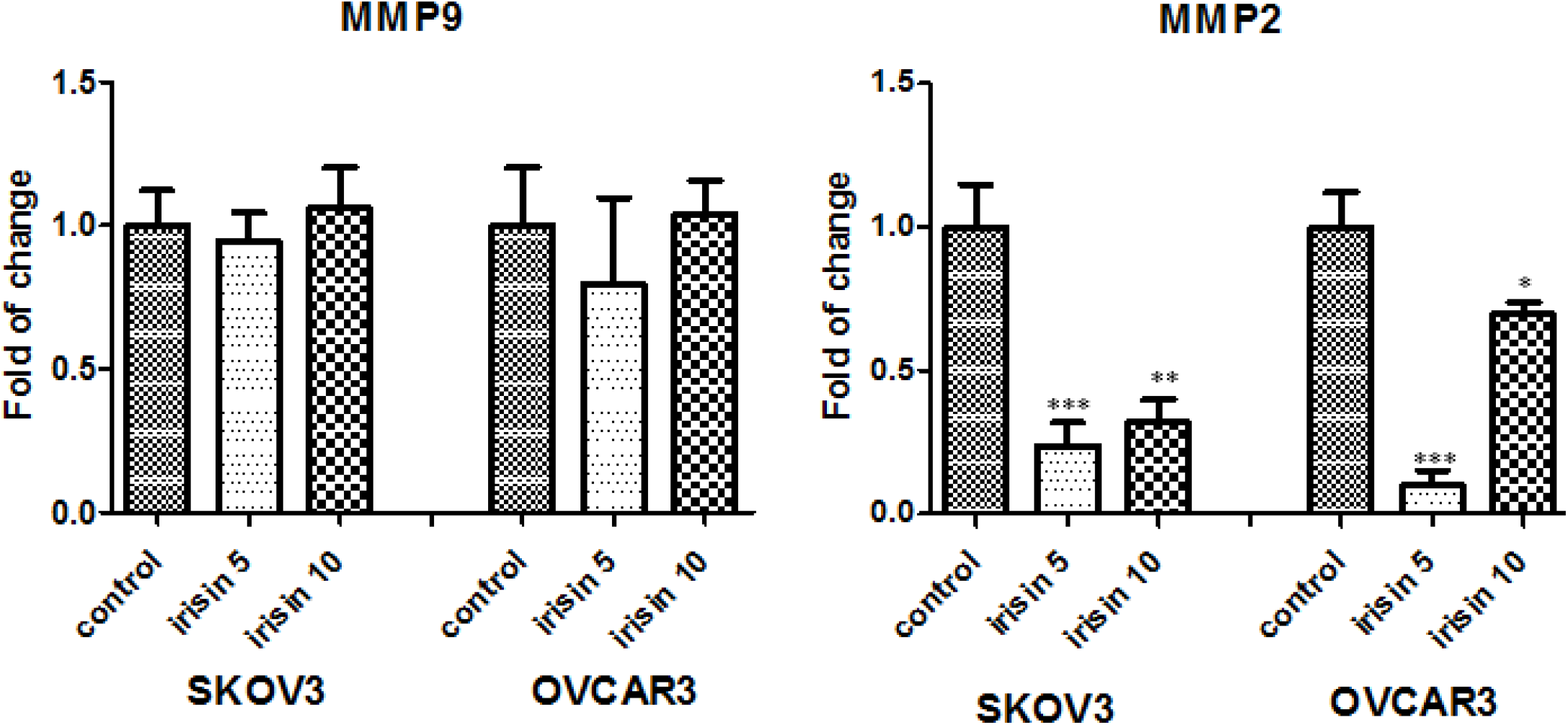
Effect of Irisin on MMP2 and MMP9 expression levels 48h after treatment with 5nM and 10nM of irisin. **Notes:** Data expressed as mean ± standard deviation; \*\**P*, 0.01; \*\*\**P*, 0.001 compared to control.

## 4. Discussion

Ovarian cancer, as the most frequent gynecologic malignancies, is a type of cancer accompanied with hypoxic microenvironment [34–36].Exercise is known to be a prominent component of cancer treatment strategies. However, the effect of exercise on tumor oxygenation, hypoxia and vascularization are not fully understood [37].

The purpose of the current study was to investigate the effects of irisin, as one of the novel exercise-related hormone, on proliferation and malignant facet of OC. We have also provided evidence about the influence of irisin on expression of HIF-1α and its target genes as plausible contributors in driving OC tumorigenesis.

We found that irisin suppressed the proliferation and viability of OC cells in a concentration and time-dependent mode. We further checked the effect of irisin on long-term survival of OC cells. Our data showed that irisin caused a significant decline in the colony-forming potential of OC.

Several complex processes including cell migration and invasion attributed in tumor dissemination, suggesting the potential therapeutic interventions required for targeting of extracellular matrix degradation [38]. As expected, our results illustrated that irisin significantly mitigated the migratory behavior and invasiveness of all three cell lines as opposed to control groups in a dose-dependent way, representing anti-metastatic trait of irisin.

Given the fact that matrix metalloproteinases (MMPs) mediates cancer cell invasion and metastasis in human malignancies including OC [38], in this study we aimed to explore whether irisin affect the mRNA expression level of MMP2 and MMP9 as two verified metastasis markers in OC. Our results determined that irisin markedly reduced the mRNA expression level of MMP2 in OVCAR3 and SKOV3 cells in a dose dependent manner. However, no significant decrease in the mRNA expression level of MMP9 was detected in surveyed cells after treatment with irisin. Nevertheless, the intricate mechanisms implicated with MMPs inhibition in OC cells need to be further studied. Pervious findings by Kong et al suggested that irisin reduced the expression of MMP2 and MMP9 in osteosarcoma cells via suppressing IL-6 [39].Since a number of studies displayed a link between the type of exercise and MMPs regulation, the effect of irisin on MMPs expression level in vitro may be different from which occurs in cancer patients [40]. So it is conceivable to explore the relationship between irisin, as an exercise-derived hormone and MMPs expression after different types of exercise activities in cancer patients.

In order to test the susceptibility of OC cells to apoptosis, we next examined the percentage of apoptosis of OC cells in the presence of 10nM irisin. Our analysis depicted that irisin significantly enhanced the percentage of apoptotic Caov4 cells as compared to control cells (P<0.008), however, apoptosis did not upsurged in other two cell lines.

More exhaustive understanding of cancer biology at the cellular and molecular levels have accentuated the HIF-1α pathway as a pivotal pathway in cancer development, for which targeting the HIF-1α pathway could be an emerging area of research [33].

HIF-1α, as the central regulator of oxygen homeostasis, orchestrates adaptive adjustments to hypoxic conditions [41, 42]. It is repeatedly reported that HIF-1α remarkably upregulates in OC or borderline tissues in relative to benign tissues [43, 44]. More specially, HIF-1α expression is known to be linked with clinicopathological aspects of OC and patients’ outcome including FIGO stage, histological subtype, metastasis and 5-year survival rate [45].Extensive studies has elucidated that molecular consequences of HIF-1α upregulation are induced expression of numerous genes linked with different aspects of cancer development, including proliferation (MYC), angiogenesis (VEGF), metabolism (PDK1, LDHA) and extracellular matrix disturbance (MMP2, MMP9) [46, 47].

In this study, we assessed whether irisin had suppressive effect on expression level of HIF-1α and its target genes in OC cells. Our real-time experiments verified that SKOV3 cells had a lower expression level of genes involved in HIF-1α pathway after exposure to irisin. Meanwhile, no considerable change was detected in OVCAR3 cells after treatment with irisin with the same concentration of irsin.

HIF-1 activity is induced by mTOR-altered metabolism and induced glycolysis via adaptation to hypoxia. The HIF-1 and HIF-2 heterodimers result in metabolic shift through responding to hypoxia [48]. Of these two heterodimers, HIF-1 is a leading component attributed to tumor metabolism resulting in overexpression of glucose transporters as well as PDK1, an enzyme which hamper pyruvate entrance to the TCA cycle [49].Qiao et al. found that malignant lymphomas represent noteworthy expression of HIF-1α. This expression is mediated through nuclear factor kappalight-chain-enhancer of activated B cells (NF-κB). Notably, radiotherapy of lymphoma exhibited enhanced NF-κB activation and high level of HIF-1α. This reveals that targeted therapy of HIF-1α combined with radiation therapy of lymphoma cells could improve treatment efficacy [50] The c-Myc proto-oncogene regarded as an eminent regulator in many cellular processes including cell cycle, cell proliferation and apoptosis. c-Myc expression and subsequently LDHA expression have been reported to be downregulated during exercise [51]. One additional striking mechanism is the impact of exercise on growth factors such as (VEGF) as a potent angiogenic stimulator. Disregulated of VEGF has been long known linked with the development of almost all tumors leading to distant metastasis, which is one of the main cause responsible for patient mortality [52, 53].

Considering the aforementioned points, exercise is a master regulator of many genes contributed to cancer development. Our encouraging results in line with previous studies suggest that irisin, as an exercise-derived hormone, may be a potential anticancer agent, warranting the need for more detailed investigations for having better insights about irisin’s effects as well as underlying mechanisms in cancer therapy.

## Abbreviations

OC: Ovarian cancer
OXPHOS: oxidative phosphorylation
HIF-1α: Hypoxia-inducible factor-1α
PDK1: Pyruvate dehydrogenase kinase 1
LDHA: Lactate dehydrogenase A
VEGF: Vascular endothelial growth factor
FBS: Fetal bovine serum
PBS: Phosphate buffer saline
MTT: 3-(4,5-Dimethylthiazole-yl)-2, 5-diphenyltetrazolium bromide
DMSO: Dimethylsulfoxide
GAPDH: Glyceraldehyde-3-phosphate dehydrogenase
MMPs: Matrix metalloproteinase
NF-ΚB: Nuclear factor kappalight- chain-enhancer of activated B

## Authors’ Contributions

MAZ and ESH conducted the research, analyzed and interpreted the data, and contributed to writing the manuscript. HHK conducted part of the research, analyzed and helped interpreting the data. HN provided research material, discussed the project, analyzed and interpreted the data. All authors read and approved the final manuscript.

## Acknowledgments

The authors are sincerely thankful to Dr Morteza Salimian for his cooperation with this work.

